# Application of novel 3D culture device for human omental adipocytes and macrophages derived from surgery

**DOI:** 10.1101/2024.09.16.613269

**Authors:** Miyuki Shimoji, Hiroya Akabori

## Abstract

Visceral adipose tissue plays a key role in the inflammation, inducing metabolic dysfunction. The culturing system of major components, adipocytes and <u>a</u>dipose <u>t</u>issue <u>m</u>acrophages (ATM) have been improved up to date, for example, the ceiling culture system, three-dimensional collagen gels and membrane mature adipocyte aggregate cultures (MAAC). Here we applied for a novel 3D culture device, cellbed of human omental adipocytes with ATM derived from surgery and presented the first morphological report. The pilot study has the limitation of resolution due to the thickness, however, the simple method would be a convenient assay to detect their morphological alteration after surgery. In addition, the combination of cell morphological observation on the cellbed and adipocytokine secretion capability would give an insight into the mechanism of adipose tissue-derived inflammation after surgery. The cellbed composed of high-grade silica glass fiver has been already provided the morphological study with the critical signaling transduction in human oral cancer research. The novel simple 3D device will prevail to spread the practical diversity in the functional clinical research.

## Introduction

Obesity-related chronic inflammation is implicated in the pathogenesis of metabolic dysfunction, including type 2 diabetes [1], and increased this adipose tissue-derived inflammation in type 2 diabetes reflects the positive association between cardiovascular diseases and diabetes [2]. Interestingly, Gletsu *et al*. have shown that adipose tissue-derived inflammatory cytokines may play, at least in part, a role in the development of postoperative inflammatory response to abdominal surgery [3]. Furthermore, the infiltration of macrophages in the adipose tissues precede the development of inflammation in obese mice, suggesting that they are crucial for adipose tissue-derived inflammation [4]. These ATMs are known to be activated [5], and amplify inflammatory responses through crosstalk with adipocytes [6]. The article of surgical material describes that obesity, anatomical location, and weight loss induce significant changes in the number of adipose tissue macrophages expressing M1 or M2 surface markers [7]. We thought that the morphological alteration with the crosstalk of adipocytes and M1 or M2-ATM as well as the change of number of M1 or M2-ATM plays an important role in adipose tissue-derived inflammation.

Next, in the culturing methods of adipocytes, the ceiling culture and MAAC for functional adipocyte culture have been established since they are not adherent at the bottom of culturing dishes [8; 9]. On the other hand, the novel 3D cell culture device “ cellbed” was composed of high pure silica glass fiber. Mukaisho KI *et al* have performed the elegant study of the morphology and the metabolism of human tongue cancer cell lines on 3D culture using cellbed (Japan Vilene Co.) [10; 11; 12; 13].

However, there is no morphological report of normal adipocytes with adipose tissue macrophages (ATM) on the cellbed of 3D cell culturing. While the transwell has the pore (0.4 *µ*m) on MAAC that supplies for oxygen and nutrition property to the floated human adipocytes. Therefore, we combined to use the cellbed and the transwell in this study. Here we present a pilot morphological study of human fresh adipocytes and ATM that were isolated from surgical omental tissues.

## Materials and Methods

### Human subjects

Samples of adipose tissue were collected from the visceral adipose tissue of patients undergoing elective gastrointestinal and biliary-pancreatic surgery at National Hospital Organization Higashi-Ohmi General Medical Center in Shiga, Japan. All study subjects received written and oral information before giving written informed consent for the use of the tissue. The studies were approved by The Ethical Review Board in National Hospital Organization Higashi-Ohmi General Medical Center, Japan.

### H&E staining of tissue and immunostaining of CD11b of tissue samples

The small species (approx. 5mm x 5mm x 3mm) were collected and fixed with 4% paraformaldehyde phosphate buffer solution (PFA: nacalai tesque #09154-85) for 48 h at 4 ºC. To remove fats, the samples were treated with 70% ethanol for a couple of days at room temperature prior to processing into paraffin blocks. The method of for immunohistochemistry (IHC) is described in [14]. The 2 or 4 *µ*m section was used for H&E staining and IHC of anti-human CD11b mAb (abcam #ab307523). The samples were observed by Nikon Eclipse Ti2 microscope carrying NIS-Elements imaging software.

### Isolation of human adipocytes and CD11b positive M1-like ATM (CD11b-ATM)

Figure 1 describes the procedure in detail. The adipose tissue was diced and digested in DMEM/F12 (nacalai tesque #08460-95) containing 0.1 % collagenase Type IV (gibco #17104019) for 1h at 37ºC. Digested material was passed through a 70 *µ*m filter (greiner bio-one #542070) and then centrifuged at 50 g x 10 min. The second phase of mature adipocytes were collected and washed with DMEM/F12 containing 10% FBS and 1% Penn/Strep [9]. CD11b-ATM was prepared by EasySep Human CD11b Positive Selection and Depletion Kit (STEMCELL technologies #ST-100-0744).

**Figure 1.**
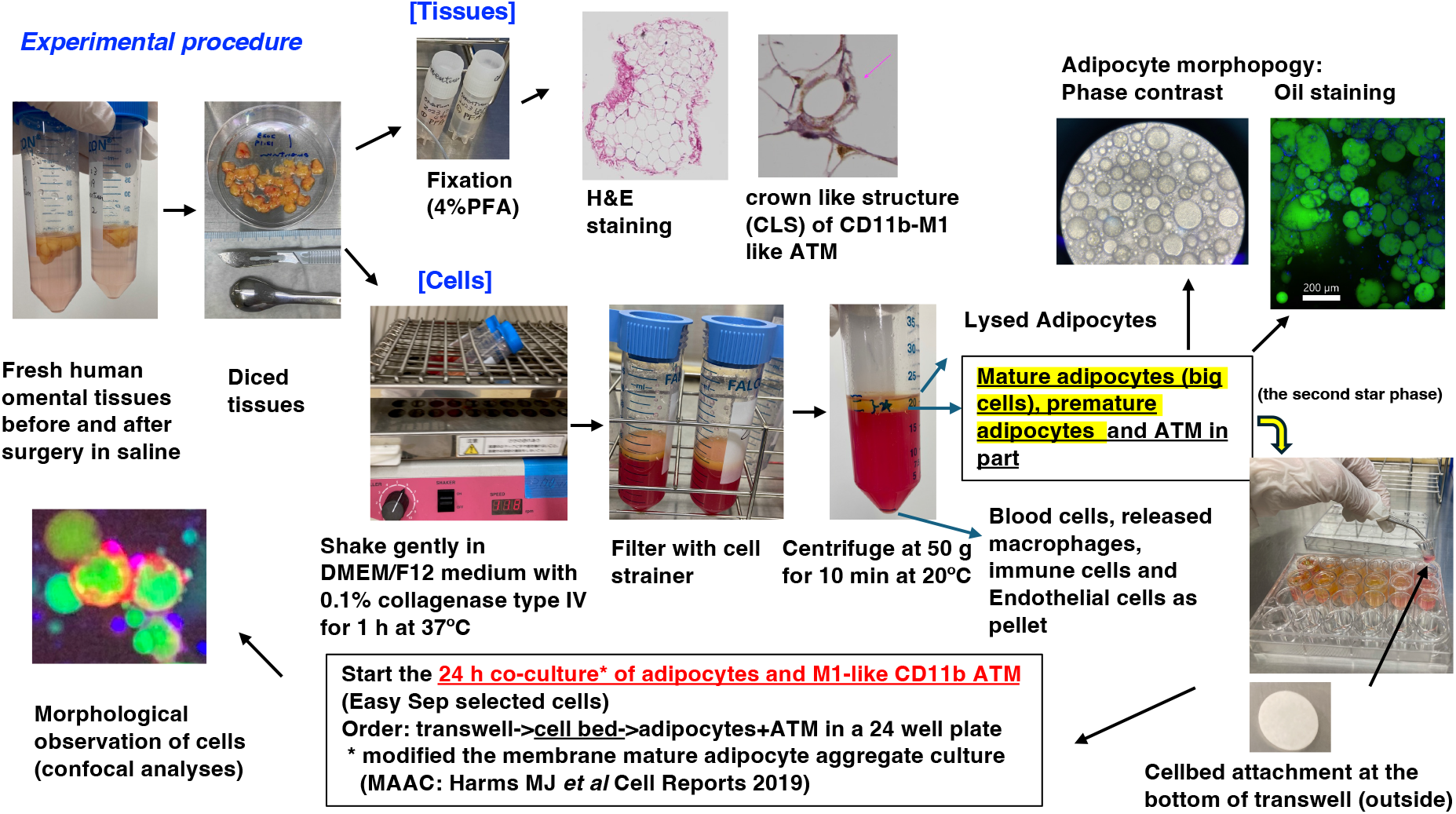
The schematic representation of experimental procedure with typical morphological results. <u>**Tissues:**</u> H&E staining of omental tissue (10x) and CLS including an adipocyte under putative cell death processing that was stained by CD11b mAb(40x) are shown in the upper pictures. <u>**Isolated cells:**</u> The picture of phase contrast observation of isolated cells after collagenese-digestion of surgical materials indicated the mixture of mature and premature adipocytes. The ratio may be dependent of the patient. The picture of bodipy staining also indicated a variety of shapes and sizes of adipocytes (the green stained cells on the upper right side). The picture of adipocytes and CD11b-ATM on the left side of bottom correspond to the round cell population at the midlle of right side in Figure 2 A. The interaction of CD11b-ATMs and adipocytes under phagocytosis was detected on the cellbed sample of 24h culturing under 5%CO2 at 37ºC in the medium of DMEM/F12 containing 1.2 nM insulin, 10%FBS and 1% Penn/Strep.

### Co-culture of reconstituted human adipocytes and CD11b-ATM

They were co-cultured by the modified MAAC culture method [9]. The suspension of human adipocytes and CD11b-ATM in DMEM/F12 (10% FBS, 1% Penn/Strep and 1.2 nM Insulin (Eli Lilly Japan)) were seeded in a 24 well plate and then set a 0.4 *µ*m pore insert (Costar-3413) with 3D cell culture device “cellbed” (Japan Vilene Co.), followed by 24 h incubation at 37ºC under 5% CO2. The adipocytes and CD11b-ATM are adherent on the surface of cellbed since the cellbed is made of high-purity silica glass fiber. The fiber creates the space to resemble for connected tissues [10].

### Imaging and Image processing

The harvested cellbed was transferred in the well of 96 well microplate (Iwaki # 3860-096). The cells on the cellbed were fixed in 2% PFA (nacalai tesque #09154-85) at room temperature for 20 min or overnight at 4ºC, permeabilized in 0.2% Triton X-100 (Sigma-Aldrich #X-100-100ML) for 20 min at room temperature and blocking in FACS buffer (10 mM HEPES, 2% FBS in PBS) for 20 min at room temperature and then stored at 4ºC until staining. The cells on the cellbed were stained with Alexa flour 647-anti-human CD11b mAb (abcam #ab307523, 1:100) for 1h at room temperature, bodipy 493/503 (Thermo Fisher Scientific #D3922, 12.5 µg/mL) for 30 min at room temperature and PureBlu Hoechst 33342 nuclear staining dye (BioRad #1351304). The cells on cell bed were washed with FACS buffer twice after each staining. The cellbed with stained cells was flipped in the well of 96 well plate to observe easily under microscope. Confocal imaging (10X) was taken on a Nikon Ti2E inverted microscope carrying Dragonfly 200 with acquisition software Fusion (Andor Oxford Instruments). The images of cells on the cellbed were imported into Imaris Software (Bioplane) for 2D and 3D visualization.

## Results and Discussion

### The first detection of human adipocytes and CD11b-ATM on cellbed of 3D cell culture

We isolated the visceral adipocytes and ATM from omental adipose tissues of surgical material, firstly. The co-culture of them for 24 h were carried out on a novel 3D culture devise” cellbed” attached with the transwell. The harvested cells on the cellbed were observed by confocal analyses. As shown in Figure 2A and 2B, the bodipy-stained mature and premature adipocytes were detected a lot (green stained round cells) and CD11b-M1 like ATM (red stained parts around the green adipocytes as indicated by a white arrow and reticular structure in part of the left side). The original omental tissue contained CLS of CD11b-ATM (upper picture of Figure 1), while CLS was detected a little in the co-cultured cells on the cellbed (data not shown). It may be consistent of the number of ATM per adipocytes [7].

**Figure 2.**
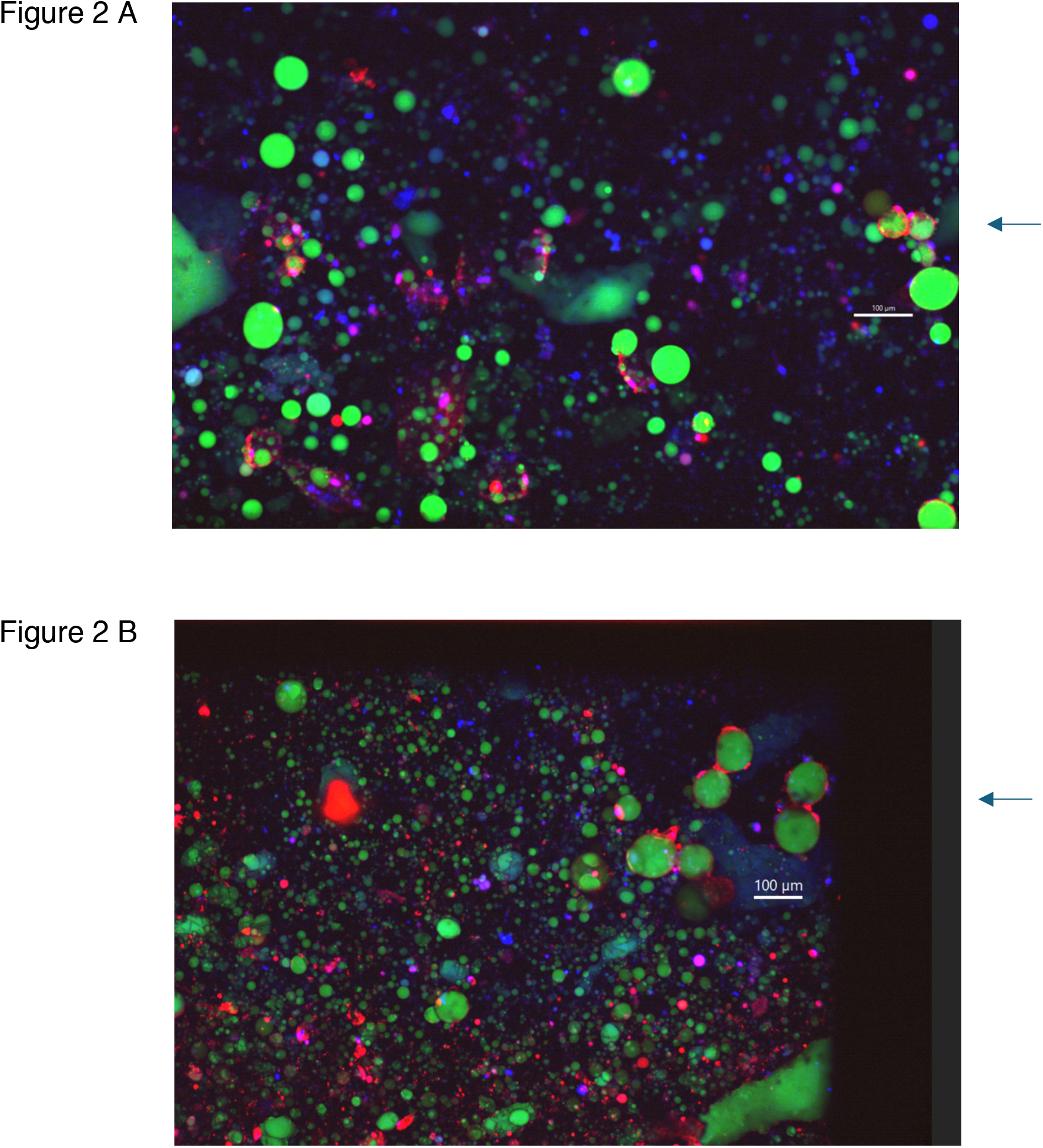
The interaction of human premature (A) and mature (B) adipocytes with CD11b-ATM. The 24 h co-culture of adipocytes and CD11b-ATM indicated CD11b-ATM embedded adipocytes under 1.2 nM insulin. It could induce the crosstalk of adipocytes and M1-like ATM, promoting the secretion of adipocytokines involved in inflammation [5]. This is currently under investigation. Scale bar: 100 *µ*m

### The limitation of resolution

We used conventional 96 microwell plates for immunostaining of cells on the cellbed in the pilot study since many numbers of cell bed were tested. Currently the method to overcome the thickness is challenged in progress e.g., using chamber slide, another microscopic analysis method etc. The 3D structural view of Figure 3 indicated the complicate localization of adipocytes and CD11b-ATM. The adipocytes are localized on either one surface of cellbed or another side, while CD11b-ATM may be extended to proliferate into the space of fibers in cellbed, resulting in the reticular structure.

**Figure 3.**
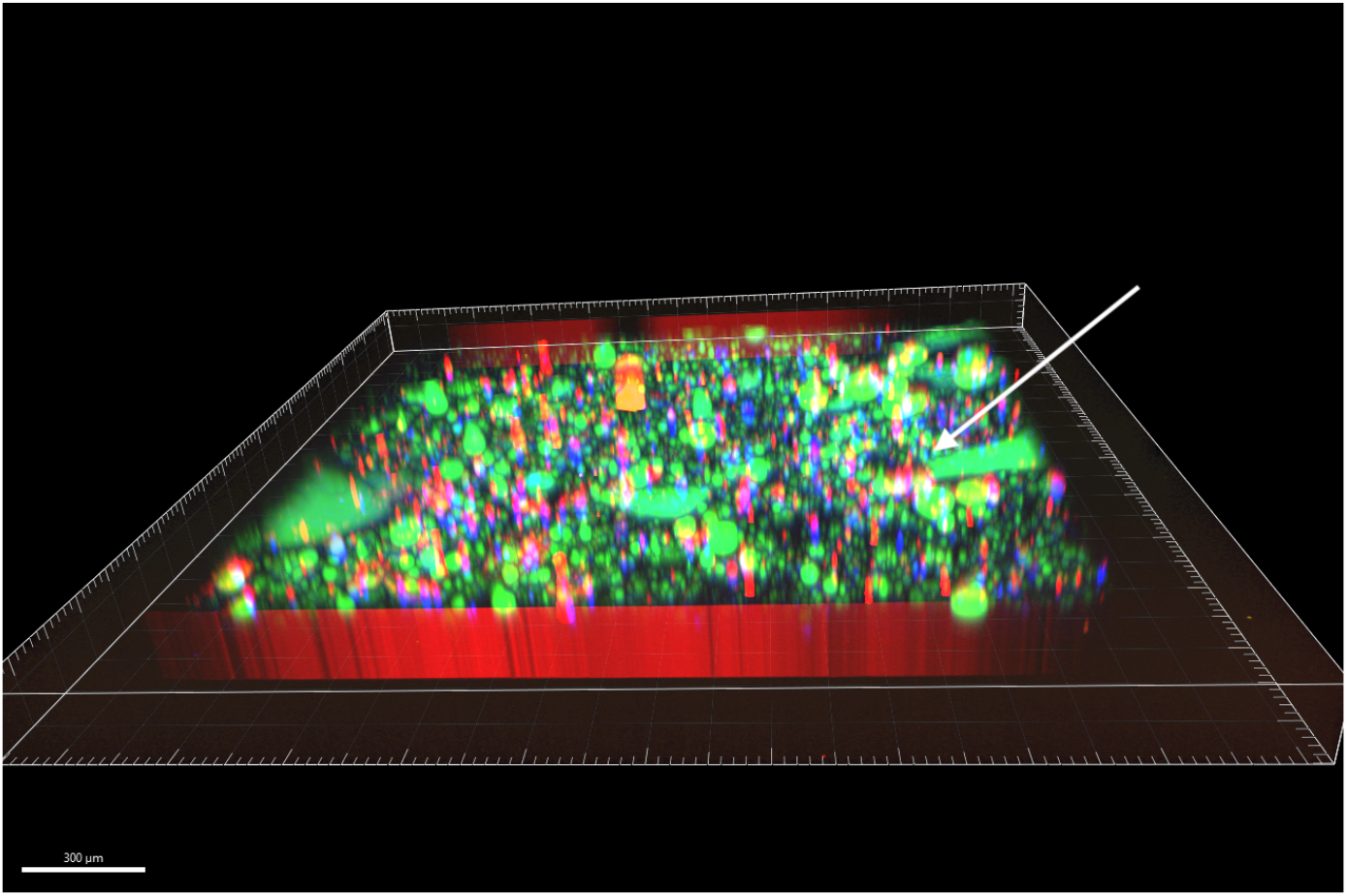
3D structural view of the cells on cellbed. It reflects on the low resolution under 10X observation, however the complicated localization of human adipocytes and CD11b-ATM suggests the crosstalks in the first pilot study. A white arrow indicates the cells corresponding to the adipocytes with CD11b-ATM of the picture on the left side in Figure 1 and the cell population near the scale bar on the right side of Figure 2 A. Scale bar: 300 *µ*m

## Conclusions

Cellbed provided the results of normal human adipocytes that interacts with ATM in this preliminary study. The limitation of resolution is caused by the thickness of cellbed. However the adipocytes and ATM on the surface of cell bed could be an easy model to calculate the functional relationship between the cellular morphology and secretion of the adipocytokines, resulting in the acute inflammation induced with visceral adipose in surgery.

## Acknowledgements

We thank Dr. Toshiyuki Watanabe (Andor Oxford Instruments), Mr. Takefumi Yamamoto (Central Research Laboratory, Shiga University of Medical Science), Ms. Arikawa, Ms. Itoh and Ms. Sawada (Department of Surgery, Shiga University of Medical Science) for technical support. We are grateful for the fruitful discussion and advice of Dr. Satoshi Murata (Cancer center of Shiga University of Medical Science Hospital), Dr. Masaji Tani (Department of Surgery, Shiga University of Medical Science), Dr. Eiji Mekata (Department of Comprehensive Surgery, Shiga University of Medical Science) and Dr. Ken-ich Mukaisho (Education Center for Medicine and Nursing, Shiga University of Medical Science, Japan).

## References

[1] A. Viardot, R.V. Lord, and K. Samaras, The effects of weight loss and gastric banding on the innate and adaptive immune system in type 2 diabetes and prediabetes. J Clin Endocrinol Metab 95 (2010) 2845–50.

[2] H. Zhang, and C. Zhang, Adipose “talks” to distant organs to regulate insulin sensitivity and vascular function. Obesity (Silver Spring) 18 (2010) 2071–6.

[3] N. Gletsu, E. Lin, J.L. Zhu, L. Khaitan, B.J. Ramshaw, P.K. Farmer, T.R. Ziegler, D.A. Papanicolaou, and C.D. Smith, Increased plasma interleukin 6 concentrations and exaggerated adipose tissue interleukin 6 content in severely obese patients after operative trauma. Surgery 140 (2006) 50–7.

[4] S.P. Weisberg, D. Hunter, R. Huber, J. Lemieux, S. Slaymaker, K. Vaddi, I. Charo, R.L. Leibel, and A.W. Ferrante, Jr., CCR2 modulates inflammatory and metabolic effects of high-fat feeding. J Clin Invest 116 (2006) 115–24.

[5] G.S. Hotamisligil, Inflammation and metabolic disorders. Nature 444 (2006) 860–7.

[6] J.M. Olefsky, and C.K. Glass, Macrophages, inflammation, and insulin resistance. Annu Rev Physiol 72 (2010) 219–46.

[7] J. Aron-Wisnewsky, J. Tordjman, C. Poitou, F. Darakhshan, D. Hugol, A. Basdevant, A. Aissat, M. Guerre-Millo, and K. Clement, Human adipose tissue macrophages: m1 and m2 cell surface markers in subcutaneous and omental depots and after weight loss. J Clin Endocrinol Metab 94 (2009) 4619–23.

[8] S. Toda, K. Uchihashi, S. Aoki, E. Sonoda, F. Yamasaki, M. Piao, A. Ootani, N. Yonemitsu, and H. Sugihara, Adipose tissue-organotypic culture system as a promising model for studying adipose tissue biology and regeneration. Organogenesis 5 (2009) 50–6.

[9] M.J. Harms, Q. Li, S. Lee, C. Zhang, B. Kull, S. Hallen, A. Thorell, I. Alexandersson, C.E. Hagberg, X.R. Peng, A. Mardinoglu, K.L. Spalding, and J. Boucher, Mature Human White Adipocytes Cultured under Membranes Maintain Identity, Function, and Can Transdifferentiate into Brown-like Adipocytes. Cell Rep 27 (2019) 213–225 e5.

[10] M. Noi, K.I. Mukaisho, S. Yoshida, S. Murakami, S. Koshinuma, T. Adachi, Y. Machida, M. Yamori, T. Nakayama, G. Yamamoto, and H. Sugihara, ERK phosphorylation functions in invadopodia formation in tongue cancer cells in a novel silicate fibre-based 3D cell culture system. Int J Oral Sci 10 (2018) 30.

[11] M. Noi, K.I. Mukaisho, S. Murakami, S. Koshinuma, Y. Machida, M. Yamori, T. Nakayama, T. Ogawa, Y. Nakata, T. Shimizu, G. Yamamoto, and H. Sugihara, Expressions of ezrin, ERK, STAT3, and AKT in tongue cancer and association with tumor characteristics and patient survival. Clin Exp Dent Res 6 (2020) 420–427.

[12] R. Ikari, K.I. Mukaisho, S. Kageyama, M. Nagasawa, S. Kubota, T. Nakayama, S. Murakami, N. Taniura, H. Tanaka, R.P. Kushima, and A. Kawauchi, Differences in the Central Energy Metabolism of Cancer Cells between Conventional 2D and Novel 3D Culture Systems. Int J Mol Sci 22 (2021).

[13] S. Murakami, H. Tanaka, T. Nakayama, N. Taniura, T. Miyake, M. Tani, R. Kushima, G. Yamamoto, H. Sugihara, and K.I. Mukaisho, Similarities and differences in metabolites of tongue cancer cells among two- and three-dimensional cultures and xenografts. Cancer Sci 112 (2021) 918–931.

[14] S. Misaki, S. Murata, M. Shimoji, T. Iwai, A.M. Sihombing, K. Aoki, Y. Takahashi, and Y. Watanabe, Enhancement of antitumor immune response by radiation therapy combined with dual immune checkpoint inhibitor in a metastatic model of HER2-positive murine tumor. Jpn J Radiol 40 (2022) 1307–1315.

